# Urine glucose levels are disordered before blood glucose levels increase in Zucker diabetic fatty rats

**DOI:** 10.1101/122283

**Authors:** Wei Yin, Weiwei Qin, Youhe Gao

## Abstract

Diabetes mellitus is a type of metabolic disease marked by hyperglycemia, and 90% of diabetes cases are type 2 diabetes mellitus. More than half of patients are not diagnosed at all or are diagnosed too late to be effectively treated, resulting in nonspecific symptoms and a long period of incubation of the disease. Pre-diabetes mellitus, also known as impaired glucose regulation, is an early warning signal of diabetes, which has long been determinated by impaired fasting glucose and impaired glucose tolerance. In this study, Zucker diabetic fatty (ZDF) rats were used to test if there were changes in urine glucose before blood glucose increases. Six 8-week-old male ZDF rats (fa/fa) and Zucker lean (ZL) rats (fa/+) were fed with Purina 5008 high-fat diet and tested for fasting blood glucose and urine glucose. After 12 weeks of feeding, the urine glucose values of the ZL rats were normal (0-10 mmol/L), but the values of the ZDF model rats increased 10 weeks before their blood glucose levels elevated. The urine glucose values of the ZDF model rats showed a state of disorder that was frequently elevated (>10 mmol/L) and occasionally normal (0-10 mmol/L). This finding may provide an easy early diagnosis. Screening for human diabetes can be considered by frequently monitoring urine glucose levels: pre-diabetes may be revealed by frequently disordered urine glucose levels over a period.

## Introduction

According to a survey by the World Health Organization, the number of global diabetic patients increased from 108 million in 1980 to 422 million in 2014, which contained 4.7% of the world's adults in 1980 and 8.5% in 2014 (W.H.O., 2016). Currently, China has the largest number of diabetic patients, approximately 114 million, with 11.6% of adult men suffering from diabetes, of which approximately 90% is type 2 diabetes mellitus; by 2040, the number may increase to 150 million (Xu et al., 2013). Without ideal treatment, diabetes may cause many complications such as cardiovascular and cerebrovascular disease, kidney, retinal and nervous chronic diseases, and various infections; even severe ketoacidosis and disability or death in the worst may occur (Danaei et al., 2014; Seuring et al., 2015).

The early symptoms of diabetes are not obvious. The disease has a long incubation period; therefore, many patients did not even realize they had developed the disease. A large part of them had contracted complications in the brain, heart, eye, kidney and other important organs, which were the main reasons for the high disability and mortality rate of diabetes, burdens both the patients and the health care system. The early diagnosis of diabetes is the prerequisite for early intervention, which is very necessary to reduce the burden (Tabák et al., 2012).

Pre-diabetes mellitus, also known as impaired glucose regulation^6^, is an intermediate state of diabetes in healthy people, which is considered a necessary stage and an early warning signal of diabetes. Impaired glucose regulation contains impaired fasting glucose and impaired glucose tolerance (Alberti and Zimmet, 1998; W.H.O., 1985). Impaired glucose tolerance refers to the special metabolic state with slightly higher postprandial blood glucose levels between healthy and diabetic patients while fasting blood glucose levels are normal. Currently, it is believed that impaired glucose regulation is an early manifestation of the pathogenetic process of diabetes, especially in type 2 diabetes mellitus. The pre-diabetes patient number in China may reach 493.4 million, which has become a serious public health problem (Yang et al., 2012).

The 2016 American Diabetes Association guidelines for the diagnosis and treatment criteria of pre-diabetes are as follows: impaired fasting glucose of fasting blood glucose from 5.6 to 6.9 mmol·L^-1^ and impaired glucose tolerance of 2 hours postprandial blood glucose from 7.8 to 11.1 mmol· L^-1^ (A.D.A., 2016). Pre-diabetes is the important transitional stage from normal glucose tolerance to diabetes. However, at the same time, pancreatic β cells of impaired glucose tolerance patients still maintain some compensatory capacity, which significantly delays or even prevents the majority of occurrence of diabetes through reasonable intervention. Therefore, rapid, convenient and effective diagnosis of pre-diabetes is critical to preventing and controlling diabetes (Bansal, 2015).

For the public, fasting blood glucose and impaired glucose tolerance testing is too complex to monitor daily at home, and the process of measuring blood glucose is not comfortable for everyone, especially before a definite diagnosis. In contrast, the measurement of urine glucose, which is simple and non-invasive, is more convenient and highly accepted, and it can be repeated more frequently at any time.

ZDF rats are universally used as the model of congenital inherited type 2 diabetes mellitus and obesity because they have hereditary characteristics of insulin receptor deletion (Zucker and Zucker, 1961; Chua et al., 1996). Male obese ZDF rats are fed a Purina 5008 high-fat diet and are characterized by hyperinsulinemia and hyperglycemia. ZDF rats develop type 2 diabetes mellitus often associated with hyperlipidemia and hypertension, which is similar to humans in biochemical metabolism, histomorphological changes and cardiovascular complications; its complications are analogous to human insulin-resistant diabetes accompanied by hypertension. The majority of patients with type 2 diabetes mellitus have this feature, making the ZDF rat model very suitable for the study of type 2 diabetes mellitus and its complications (Durham and Truett, 2006). Corresponding to ZDF rats, ZL rats are non-obese and non-diabetic animals with diminished stature, whose weight differs from ZDF rats over time, and they are used as the normal control for ZDF rats.

In this study, ZDF rats were used to study the change of urine before type 2 diabetes mellitus diagnosis to find a convenient, non-invasive early indicator of impaired glucose regulation.

## Materials and Methods

### Animal Experiments

This study consisted of six male ZDF rats (fa/fa) and six male ZL rats (fa/+). Animals were received from Beijing Vital River Laboratory Animal Technology Co., Ltd. (Beijing, USA) at 8 weeks of age and were individually caged to allow for individual measurements of food consumption. All animals were fed Purina 5008 rat chow (protein =23%, carbohydrate =58.5%, fat =6.5%, fiber =4%, and ash =8% by weight). All rats were maintained under stringent environmental conditions that included strict adherence to 12-hour light/dark cycles. All animal manipulations and care procedures were carried out between 1.5 hours and 3.5 hours after lights on. Animals were fed ad libitum from 8 weeks until 24 weeks of age and then sacrificed by aortic exsanguination. All protocols adhered to the “Guide for the Care and Use of Laboratory Animals” and were approved by our institution’s Institutional Animal Care and Use Committee.

### Experimental design

Once a week, rats were placed individually in metabolic cages fasting for 12 hours, and urine was collected. Then, urine volume and body weights were recorded, and blood was sampled from the retrobulbar venous plexus for analysis of fasting blood glucose measured with glucometer (ACCU-CHEK^®^, Roche Diabetes Care, Inc., Indianapolis, Indiana). Urine glucose concentrations were analyzed with a urine glucose assay kit (Oxidase method, Beijing Applygen Technologies Inc., Beijing, China).

## Results

### Metabolic parameters

During the treatment period of 16 weeks (age of the rats: 8–24 weeks), there were no spontaneous deaths in all animals. ZL rats showed good growth, flexible activities and healthy coat color. ZDF rats had increased feed and water intake, increased urine output and excrement and dull coat color. ZDF rats showed a significant increase in body weight (P < 0.05), which increased rapidly from 8 to 12 weeks and slowly after 12 weeks. ZL rats showed a continuous and stable increase. The body weight changes of ZDF rats and ZL rats are shown in Fig. 1.

**Fig. 1.**
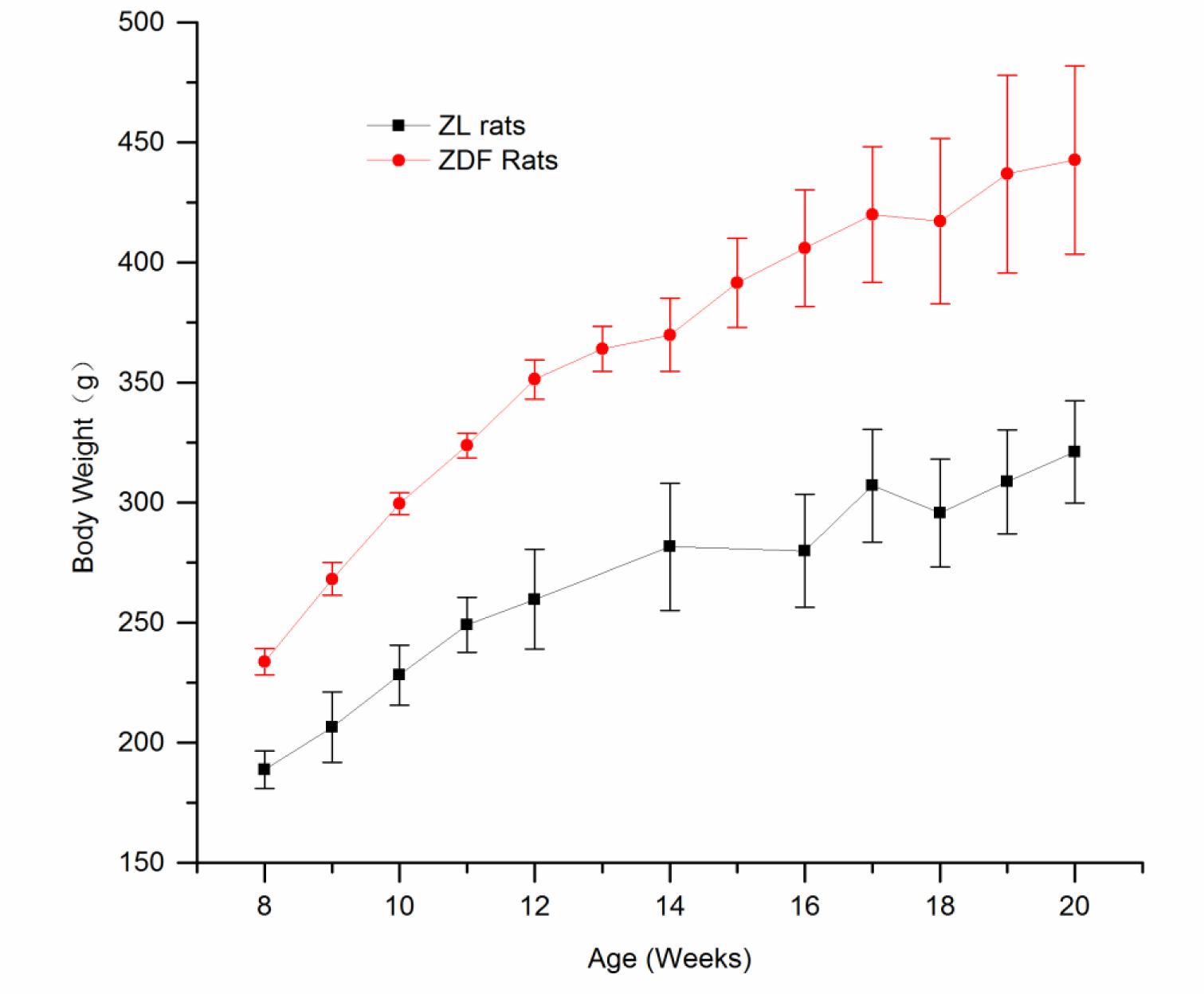
The body weight changes of ZDF rats and ZL rats. Body weight of ZDF rats (red circles) and ZL rats (black squares) (n =6, mean±SEM, fed conditions). Symbols represent means, and error bars represent one standard deviation. ZDF rats were significantly different from ZL rats (P<0.05).

### Glucose changes

The fasting blood glucose level of ZDF rats increased obviously from 16 weeks and was significantly higher than ZL rats (P <0.05), as shown in Figure 2. The urine glucose concentration of ZL rats was always between 0-10 mmol·L^-1^. The concentration in ZDF rats was occasionally lower than 10 mmol·L^-1^, but most of the time, it was higher than 10 mmol·L^-1^. The concentration always higher than 10 mmol·L^-1^ when the fasting blood glucose level turned to rapid elevation phase, as shown in Fig.2.

**Fig. 2.**
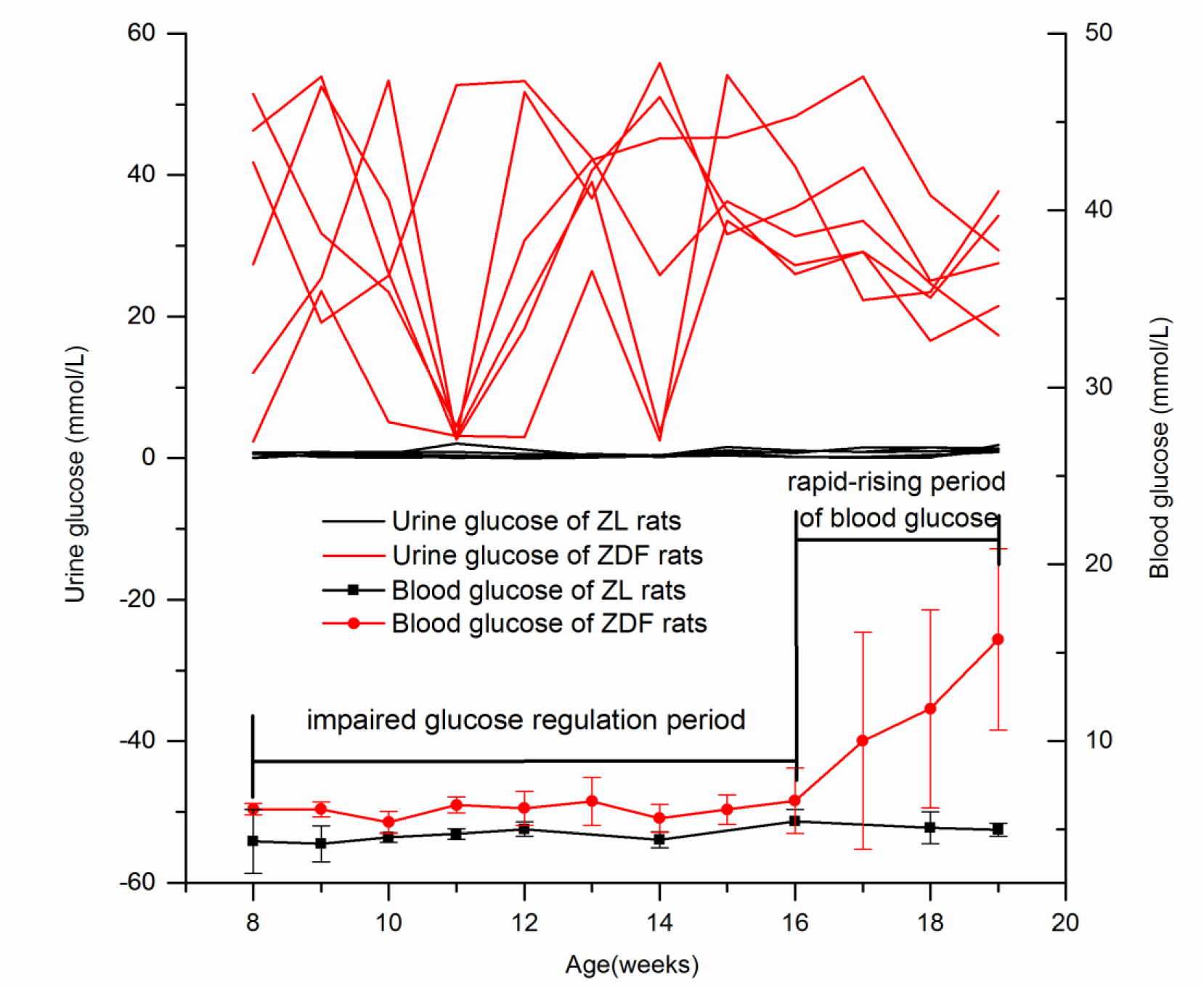
Fasting blood glucose and urine glucose changes of ZDF rats and ZL rats. Fasting blood glucose of ZDF rats (red circles) and ZL rats (black squares) (n =6, mean ± SEM, fed conditions). Symbols represent means, and error bars represent one standard deviation. ZDF rats were significantly different from ZL rats after week 16 (P < 0.05). Urine glucose concentration of ZDF rats (red lines) and ZL rats (black lines) (n =6, fed conditions). ZDF rats were disordered and significantly different from ZL rats, which were always low.

## Discussion

The kidney is the main excretory organ; most of the glucose in glomerular filtrate will be reabsorbed back into the blood by renal tubules in healthy individuals. Only a very small amount of glucose remains in urine, which cannot be detected in a urine test. However, the glucose reabsorption by proximal tubules is limited. When blood glucose concentration is more than 8.96∼10.08 mmol/L, epithelial cells of proximal tubule reaches the limit of glucose absorption and thus glucose cannot be fully reabsorbed and excreted into urine, which is called glycosuria. The lowest blood glucose concentration at which glucose begins to appear in urine is known as renal threshold for glucose (RT_G_). When the blood glucose concentration exceeds the RT_G_, urine glucose begins to appear (Polidori et al., 2013; Polidori et al., 2013).

Although the blood glucose level of ZDF rats is higher than ZL rats, it is still in the normal range and no higher than the RT_G_ of ZDF rats (Liang et al., 2012). Before crossing the key point of elevated blood glucose, the increasing urine blood level may be due to impaired glucose regulation, which caused short-term and uncaptured changes when blood glucose level exceeds the RT_G_. As urine glucose is monitored over a period, unlike in blood that is monitored at a point of time, urine glucose is easily captured, and the disorder of urine glucose level can be observed. Previous studies have produced similar findings on ZDF rats (Hempe et al., 2012) and GK diabetic Rats (Ueta et al., 2005) regarding urine glucose disorder before blood glucose goes beyond the normal range and regarding how it may be a valuable early warning sign.

Based on the results of this study, during early impaired glucose regulation, urine glucose disorder may be caused by occasional blood glucose increase over the RT_G_ at some points. When glucose is filtered into the urine, glycosuria begins to appear. Because of the ability of blood to maintain steady state is strong, blood glucose is normal most of the time, especially after 12 hours of fasting. During impaired glucose regulation, blood glucose increases, which reflects a status of a certain time point and may be difficult to capture. However, since urine can accumulate for a period, the glucose increase can be captured easily in urine. Therefore, in the pre-diabetes period, urine can promptly reflect the appearance of impaired glucose regulation by the disorder of urine glucose levels.

It is likely that the more serious impaired glucose regulation becomes, the greater the frequency of abnormal urine glucose levels tend to be. When the fasting blood glucose concentration increases, urine glucose levels stay high. This phenomenon provides a new thought for the early warning of type 2 diabetes mellitus, suggesting that human diabetes screening can be carried out by frequent urine glucose monitoring. If the above findings remain true in humans, the frequency of high urine glucose level may be an indicator to impaired glucose regulation, which can be used for simple, noninvasive and effective home self-monitoring. Intelligent closestools, which can provide easy frequent urine glucose testing at home, may help to identify pre-diabetes people.

## Statistical analysis

All statistical analyses were carried out by using SPSS 22.0 software (IBM). Results are expressed as mean±s.d. from the experiments that repeated at least three times, Student’s t-test or one-way ANOVA analysis were used to compare differences between groups. P-values <0.05 were considered statistically significant.

## Author contributions

Designed the experiments: Youhe Gao, Performed the experiments: Wei Yin and Weiwei Qin, analyzed the data: Wei Yin, Wrote the paper: Youhe Gao and Wei Yin.

## References

Alberti, K.G.M.M. and Zimmet, P. Z. (1998). Definition, diagnosis and classification of diabetes mellitus and its complications. Part 1: diagnosis and classification of diabetes mellitus. Provisional report of a WHO consultation. Diabetic medicine. 15, 7539–553.

American Diabetes Association. (2016). Standards of medical care in diabetes—2016. Diabetes care. 39, S16.

Bansal, N. (2015). Prediabetes diagnosis and treatment: A review. World journal of diabetes. 6, 296.

Chua, S. C.,Chung, W. K., Wu-Peng, X. S., Zhang, Y., Liu, S. M., Tartaglia, L. and Leibel, R. L. (1996). Phenotypes of mouse diabetes and rat fatty due to mutations in the OB (leptin) receptor. Science. 271, 994–996.

Danaei, G., Lu, Y., Singh, G. M., Carnahan, E., Stevens, G. A., Cowan, M. J., Farzadfar, F., Lin, J. K., Finucane, M. M., Rao, M. et al. (2014). Cardiovascular disease, chronic kidney disease, and diabetes mortality burden of cardiometabolic risk factors from 1980 to 2010: a comparative risk assessment. The Lancet Diabetes and Endocrinology. 2, 634–647.

Durham, H. A. and Truett, G. E. (2006). Development of insulin resistance and hy-perphagia in Zucker fatty rats. American Journal of Physiology - Regulatory, Integrative and Comparative Physiology. 290, 652–658.

Hempe, J., Elvert, R., Schmidts, H. L., Kramer, W. and Herling, A. W. (2012). Appropriateness of the zucker diabetic fatty rat as a model for diabetic microvascular late complications. Laboratory Animals. 46, 32–9.

Liang, Y., Arakawa, K., Ueta, K., Matsushita, Y., Kuriyama, C., Martin, T., Du, F. Y., Liu, Y., Xu, J., Conway, B., et al. (2012). Effect of canagliflozin on renal threshold for glucose, glycemia, and body weight in normal and diabetic animal models. Plos One. 7, e30555–30555.

Polidori, D., Sha, S., Ghosh, A., Plum-Mörschel, L., Heise, T., and Rothenberg, P. (2013). Validation of a novel method for determining the renal threshold for glucose excretion in untreated and canagliflozin-treated subjects with type 2 diabetes mellitus. The Journal of Clinical Endocrinology & Metabolism. 98, E867–E871.

Polidori, D., Sha, S., Mudaliar, S., Ciaraldi, T. P., Ghosh, A., Vaccaro, N., Farrell, K., Rothenberg, P. and Henry, R. R. (2013). Canagliflozin lowers postprandial glucose and insulin by delaying intestinal glucose absorption in addition to increasing urinary glucose excretion. Diabetes care. 36, 2154–2161.

Seuring, T., Archangelidi, O. and Suhrcke, M. (2015). The economic costs of type 2 diabetes: a global systematic review. Pharmacoeconomics. 33, 811–831.

Tabák, A. G., Herder, C., Rathmann, W., Brunner, E. J. and Kivimäki M. (2012). Prediabetes: a high-risk state for diabetes development. The Lancet. 379, 2279–2290.

Ueta, K., Ishihara, T., Matsumoto, Y., Oku, Q., Nawano, M., Fujita, T. et al. Long-term treatment with the Na+-glucose cotransporter inhibitor T-1095 causes sustained improvement in hyperglycemia and prevents diabetic neuropathy in Goto-Kakizaki Rats. (2005). Life Sciences. 76, 2655–2668.

World Health Organization. (1985). Diabetes Mellitus: Report of a WHO Study Group. Geneva: WHO, Technical Report Series. 727.

World Health Organization. (2016). Global report on diabetes. Geneva, 2016. Available from: http://apps.who.int/iris/handle/10665/204871.

Xu, Y., Wang, L., He, J., Bi, Y., Li, M., Wang, T., Wang, T., Jiang, Y., Dai, M., Lu, J. et al. (2013). Prevalence and Control of Diabetes in Chinese Adults. JAMA. 310, 948–59.

Yang, W. Y., Zhao, W. H., Xiao, J. Z., Li, R., Zhang, P., Kissimova-Skarbek, K., Schneider, E., Jia, W. P., Ji, L. N., Guo, X. H. et al. (2012). Medical care and payment for diabetes in China: enormous threat and great opportunity. PloS one. 7, e39513.

Zucker, L. M. and Zucker, T. F. (1961). Fatty, a new mutation in the rat. Journal of Heredity. 52, 275–278.

